# Development and on-site evaluation of an easy-to-perform and low-cost food pathogen diagnostic workflow for low-resource communities

**DOI:** 10.1101/2020.05.29.122994

**Authors:** Michael Glenn Mason, José Ramón Botella

**Affiliations:** Plant Genetic Engineering Laboratory, School of Agriculture and Food Sciences, The University of Queensland, Australia

## Abstract

Food-borne diseases are a leading cause of illness and death in many developing countries and thus, there is a real need to develop affordable and practical technologies that can help improve food safety in these countries. The ability to efficiently identify food pathogens is essential to allow national regulatory authorities to monitor food quality and implement safety protocols. In this study, we have developed a simple, low-cost ($0.76 (USD)) complete food pathogen diagnostic workflow ideally suited for deployment in low-resource environments that uses a simple four step process (sample enrichment, cell lysis, DNA amplification, and naked-eye readout). The minimal number of steps and equipment involved in our diagnostic workflow, as well as the simplicity of the yes/no flocculation readout, allows non-technical personnel to perform and interpret the assay. To evaluate the system’s performance, we tested the entire system on fresh produce samples collected from local farms and markets in Cambodia for the presence of the *E. coli O157* O-antigen polymerase, *wzy*. Although this was a proof-of-concept study, our system successfully revealed a clear correlation between the origin and condition of the produce collected and their likelihood of contamination. In conclusion, we believe that our easy-to-perform diagnostic system can have a significant impact on improving food quality and human health if adopted by regulatory authorities in developing countries due to the assay’s simplicity, affordability, and adaptability.

## Introduction

Over the last 20 years, there has been a significant increase in the incidence of food-borne diseases worldwide [1]. The risk of infection by a food-borne disease is significantly higher in developing countries due to a combination of factors including access to clean water and bathroom facilities, poor hygiene education, inadequate food production and storage practices, and either insufficient food safety legislation or poor implementation of existing legislation [2]. Most food-borne disease infections in these countries result from the consumption of perishable foods sold in informal markets [3] and, as such, food-borne diseases have become a leading cause of illness and death in developing countries [4, 5]. Hence, there is a need for low-cost and simple technologies that can help health authorities to monitor and enforce adequate levels of food safety. In addition to the health benefits, increased food safety standards would likely have economic benefits as a result of increased demand for fresh produce exports [3].

Shiga toxin-producing *Escherichia coli* (STEC) are a class of *E. coli* responsible for many food-borne outbreaks and sporadic cases of gastrointestinal illness, with a range of symptoms including haemorrhagic colitis (stomach cramping and bloody diarrhoea) and the potentially fatal hemolytic-uremic syndrome (break down of red blood cells, kidney failure, reduction in platelet cells) [6]. In the United States alone, STEC strains are estimated to cause approximately 176,000 illnesses, 2,400 hospitalizations, and 20 deaths each year [7]. Of all the STEC strains, the O157:H7 serotype is the most common cause of foodborne illness outbreaks worldwide [8] and is the responsible for all of the deaths by STEC strains in the US [7]. Infection by *E. coli O157:H7* is commonly thought to be associated with the consumption of undercooked meat however, infection can also occur through the consumption of fresh fruit and vegetables [7, 9, 10]. For example, outbreaks of *E. coli O157:H7* in the United States have occurred due consumption of a variety of fresh produce including onions, bagged spinach, lettuce and unpasteurized apple juice [10–12]. Produce and water supplies used for irrigation or consumption can become contaminated by coming in contact with trace amounts of fecal waste from farm animals, which can be carriers of *E. coli O157:H7* but are themselves unaffected by it [6, 13].

*E. coli O157:H7*’s low infective dose makes it a highly effective and dangerous human pathogen as it can still cause illness even when the initial source of contamination has been significantly diluted [14]. Thus, countries typically adopt a 0 CFU limit of *E. coli O157:H7* or other STEC strains in food [15]. The standard method of identifying *E. coli O157:H7* is an involved process that requires a trained microbiologist and takes days to complete [16, 17]. While this method is well established and highly reliable, it is not always practical, especially for developing countries who have a limited number of trained personnel and testing facilities. DNA amplification-based systems can provide an alternative to detect food borne pathogens as they are low-cost, fast and relatively simple to perform [17]. Polymerase chain reaction (PCR) is the conventional DNA amplification-based technique for rapid identification of DNA/RNA biomarkers [18] however, PCR-based methods require a relatively expensive thermocycler and hence are not ideally suitable for low resource environments. Isothermal DNA amplification methods, including Loop-mediated isothermal amplification (LAMP) [19], may potentially overcome equipment costs as they enable DNA amplification to be performed in a low-cost, single temperature heating device.

We have developed a flocculation-based DNA amplification readout that enables the identification of the presence of amplification even by inexperienced users [20]. Thus, in this study, we sought to build upon the advantages of the isothermal DNA amplification and our flocculation readout technique to create a complete food pathogen diagnostic system tailored for developing countries. We have optimized the entire process, from food sampling to pathogen detection, aiming to simplify each step, reduce costs and ensure safety. In a proof-of-concept study, we used our diagnostic workflow to detect *E. coli* O157 in Cambodian fresh produce and demonstrated that our system is capable of rapidly screening fresh produce for the presence of food pathogens in low-resource environments.

## Results

### Sample processing and bacterial enrichment

In modern food microbiology laboratories, a stomacher machine is typically employed to macerate tissue in growth media prior to microbial enrichment [21]. This equipment is expensive and, in the hands of poorly trained personnel, can be a source of cross-contamination resulting in false positives and misleading results. We therefore developed an equipment free version of a stomacher machine in which samples are collected and placed in individual tubes containing growth media and 3-4 metal ball bearings (S1A Fig). Vigorous shaking of the tubes macerates the tissue and aids the release food pathogens into the media which can then be enriched in an overnight incubation without the need to open the tube, minimizing the risk of cross-contamination.

### Rapid bacterial DNA extraction from fresh produce enrichment cultures

A critical part of any food pathogen molecular diagnostic is the ability to reliably, cheaply and safely extract amplifiable DNA. As the infective dose of *E. coli O157:H7* is as low as a few hundred cells [7, 14, 22], an initial enrichment step in buffered peptone media for 16-24 hours is required before detection. We reasoned that the cheapest and safest method to extract microbial DNA from enriched samples is to heat the entire culture to 95°C for 30 minutes as this will kill the bacteria in the sample and lyse the *E. coli*, which can be performed without opening the tube. To test this hypothesis, we infected Brussel sprout leaves with *E. coli O157:H7*, and subsequently processed the leaf samples as described above. Tubes containing macerated leaf tissue in buffered peptone water were incubated at 41.5°C for 24 hours and then heated at 95°C for 30 minutes. Samples from the boiled culture failed to produce bacterial growth on buffered peptone agar plates, indicating that the heat treatment had killed the bacterial population and the tube was safe to open.

Direct addition of 1, 2 or 4 μl of the boiled culture into LAMP reactions did not result in amplification (S1B Fig) indicating that amplification inhibitors are present in the boiled extract. However, a 25-fold dilution of the boiled extract with water prior to addition to the LAMP reaction resulted in strong amplification (S1B Fig).

### LAMP primer development

LAMP primers were developed against a number of *E. coli O157:H7* genes (S1 Table), including the shiga toxin genes *stx1* and *stx2* [23], the O-antigen polymerase *wzy* [24, 25], the O157-specific biosynthesis gene, *rfbE* [26], and the H7 flagella antigen *fliC* [27]. The primers were used in LAMP reactions to detect the presence of *E. coli O157:H7* in cultures generated from artificially inoculated alfalfa sprouts in three independent experiments. The *stx2-2* primer set was unreliable as it failed to amplify a product in two of the three experiments (Fig 1A) while the *fliC*, *stx2*-2 and *rfbE* primer sets produced amplification products in one of the two non-inoculated control samples indicating that they are prone to false positives (Fig 1B). Based on these results, only the *wzy-1* and *stx2-1* primer sets were found to give reliable amplification of *E. coli O157:H7*.

**Figure 1.**
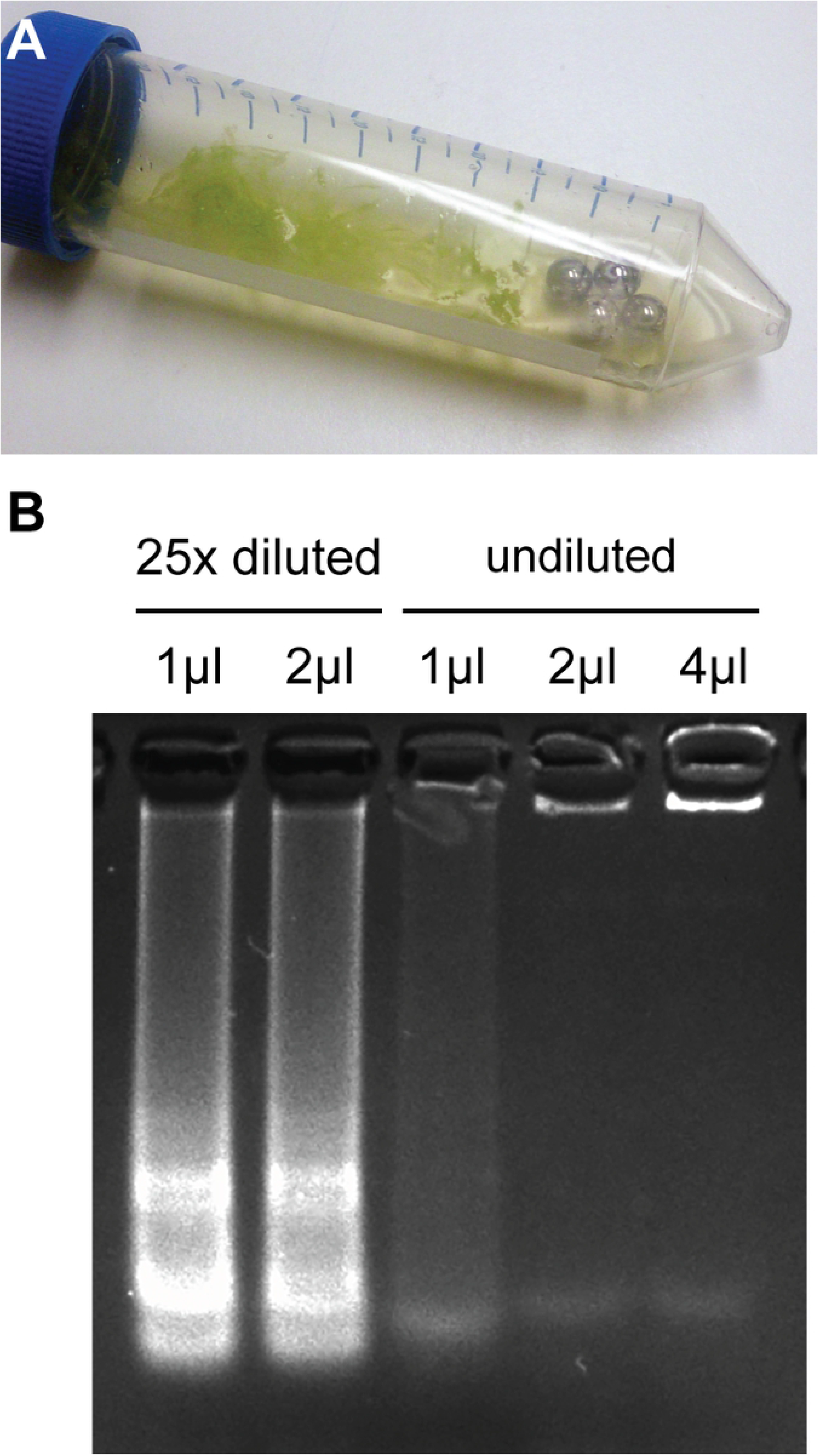
Identification of specific and reliable LAMP primers. (A) LAMP primer sets designed for *fliC-1*, *wzy-1*, *stx2-1*, *stx2-2*, *rfbE-1* were tested for their ability to detect *E. coli O157:H7* in three artificially inoculated Alfalfa sprout samples. (B) The same sets of LAMP primers were used in two uninoculated Alfalfa sprout control samples.

To further test the specificity of the *wzy-1* and *stx2-1* primer sets, we examined their ability to differentiate between *E. coli O157:H7* and *Salmonella enterica*, a genetically similar pathogenic species to *E. coli* [28]. Alfalfa sprouts were inoculated with either 8 or 80 colony forming units (CFU) of *Salmonella enterica* or 1 CFU of *E. coli O157:H7* and cultured enriched overnight. LAMP primers designed against the *Salmonella* invasion protein, *invA* (Table S1), were able to specifically identify the *Salmonella* infected alfalfa sprouts while no amplification products were detected in the *E. coli O157:H7* inoculated samples and non-inoculated controls (Fig 2A). The *wzy-1* or the *stx2-1* primer sets produced strong amplifications in the *E. coli O157:H7* infected samples but not in the *Salmonella* infected samples or the non-inoculated controls (Fig 2B).

**Figure 2.**
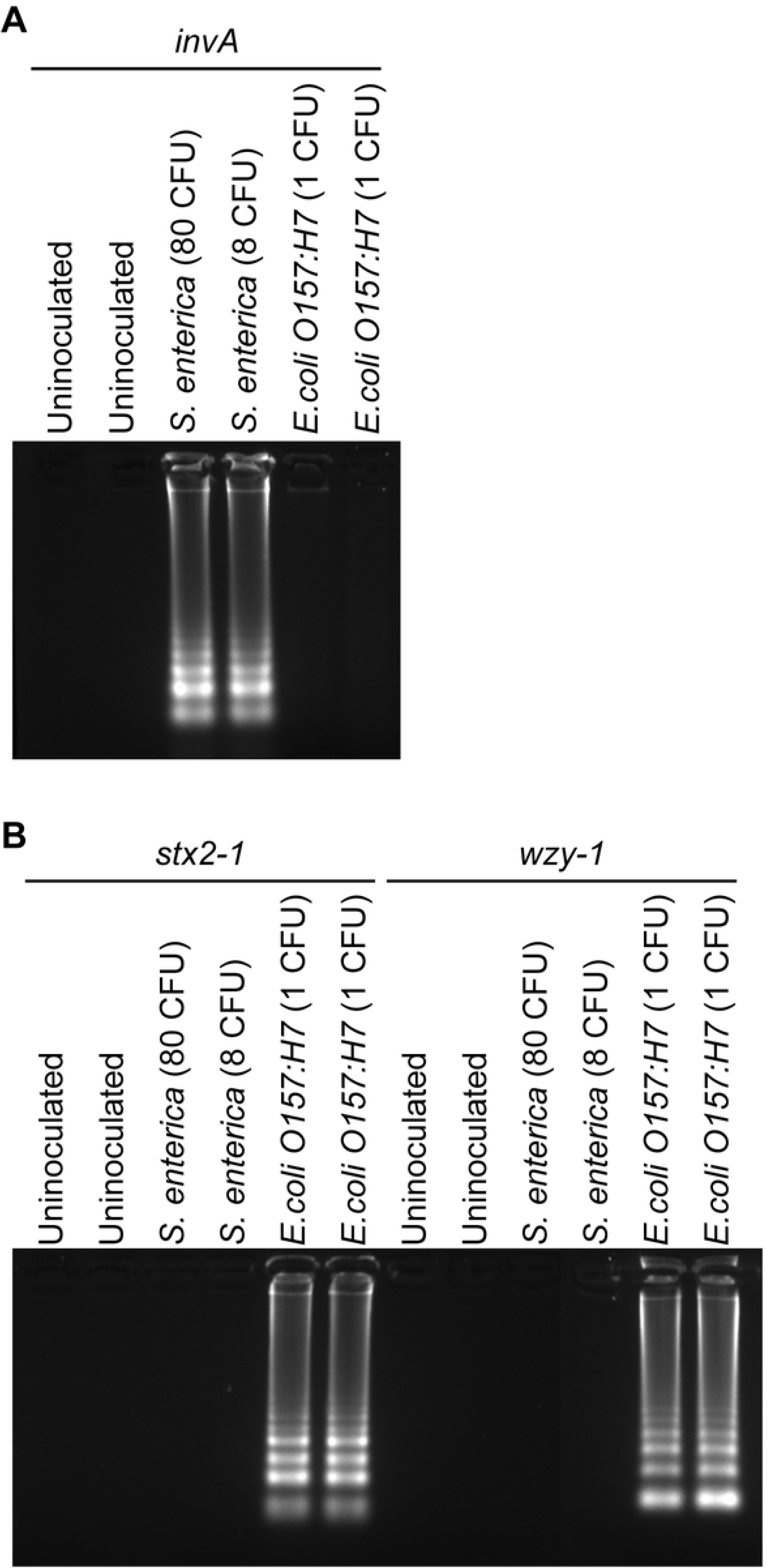
Primers *wzy-1* and *stx2-1* are specific for *E. coli O157:H7*. (A) LAMP reactions using primers targeting the *Salmonella invA* gene were performed on alfalfa sprout samples inoculated with either 8 or 80 CFU of *S. entrerica serovar enteritidis* or 1 CFU of *E. coli O157:H7*, non-inoculated sprouts were used as controls. (B) LAMP reactions using primers *wzy-1* and *stx2-1* were performed on the samples described above.

### Stabilization of LAMP reactions to allow room temperature transport

Procurement of amplification reagents in many developing countries is problematic largely due to erratic power supplies and inadequate cold storage [29, 30]. Freeze drying of reactions could facilitate room temperature transport to, and storage at remote locations. Trehalose is commonly used as a stabilizer of biomolecules during the freeze-drying process [31, 32], however our initial attempts to freeze dry a complete LAMP reaction failed to produce an amplification product upon reconstitution with water either in the presence or absence of trehalose (Fig 3A). Further tests revealed that the presence of betaine, an essential component of the LAMP reaction, negatively affected the activity of the rehydrated reaction (Fig 3A). Therefore, the LAMP reactions were freeze-dried in the absence of betaine and subsequently resuspended in a solution containing betaine at the concentration required for the reaction. To study whether the freeze drying process can stabilize the LAMP reactions for extended periods of time, the dried reactions were left at room temperature (22 - 24°C) for 4, 8 or 21 days before being rehydrated and used to amplify purified *E. coli O157:H7* genomic DNA. Strong amplifications were observed at all three time points with no observable loss of activity over time (Fig 3B).

**Figure 3.**
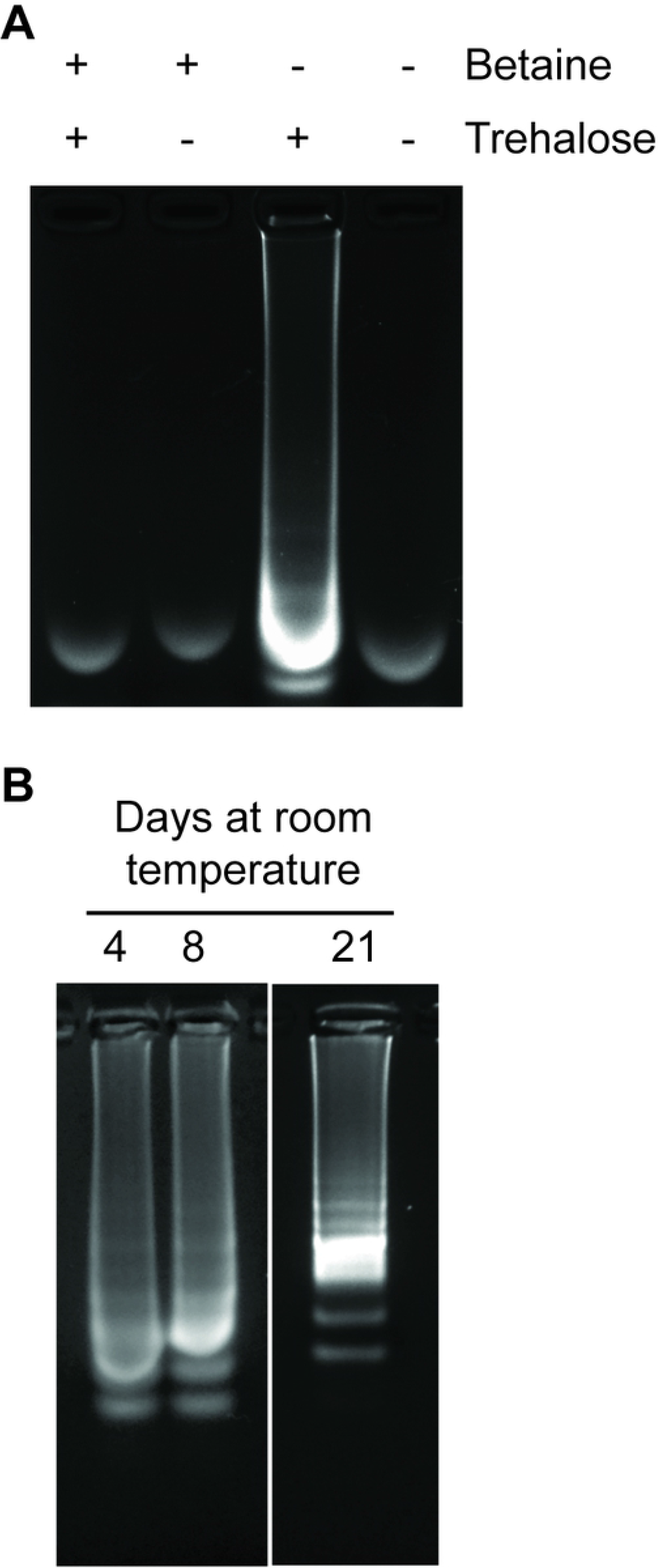
Optimization of freeze-drying conditions. (A) LAMP reactions containing all necessary components, including primers were freeze-dried overnight in the presence (+) or absence (-) of 5% (w/v) trehalose or 0.8 M betaine. Samples processed without betaine were subsequently rehydrated in betaine solution (0.8 M final concentration) prior to amplification using 2 ng of purified *E. coli O157:H7* genomic DNA as template. (B) Samples freeze-dried with trehalose but without betaine were left at room temperature for 4, 8 and 21 days prior to rehydration with betaine solution and amplification using 2 ng of purified *E. coli O157:H7* DNA as template.

### Complete diagnostic workflow and proof of concept testing in Cambodia

The individual components developed in this study were combined to create a complete diagnostic workflow suitable for testing fresh produce for *E. coli O157* contamination (Fig 4). To test the full food pathogen diagnostic system in our laboratory, 1 g of Brussel sprout leaves inoculated with *E. coli O157:H7*, were placed in a 50 ml Falcon tube containing three metal ball bearings and 10 ml of buffered peptone. Non-inoculated leaves were used as negative controls. After shaking to disrupt the tissue, tubes were incubated at 41.5°C overnight to allow enrichment and boiled at 95°C for 30 mins to kill and lyse the bacteria. Freeze dried LAMP reactions, containing *wzy-1* primers, were reconstituted with betaine solution (0.8 M betaine final concentration) and 1 μl of the enriched cultures diluted 15-fold in water added before incubating at 63°C for 50 minutes. The amplification reactions were examined by electrophoresis (Fig 5A) and by the addition of a flocculation solution [20] (Fig 5B). No amplification was detected in the water controls or the non-inoculated samples however, a strong amplification was observed in the *E. coli O157:H7* inoculated sample (Fig 5A). The addition of flocculation solution mirrored the results observed on the agarose gel that is, *E. coli* inoculated samples gave a positive flocculation result in which the particles in solution clumped together and rapidly settled to the bottom of the tube and leaving a transparent liquid phase (Fig 5B). In contrast, the water controls and the non-inoculated samples showed a negative flocculation reaction in which the black particles remained suspended.

**Figure 4.**
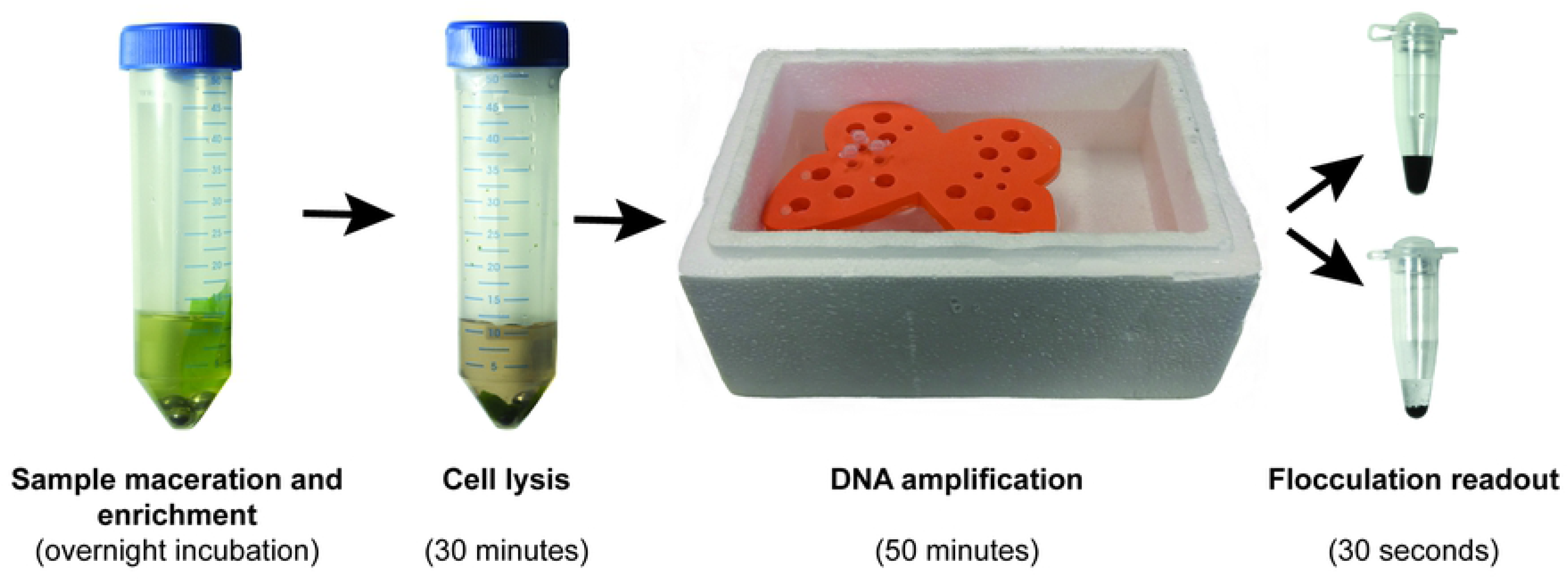
Overview of the complete food pathogen detection system. Sample maceration and enrichment involves shaking fresh produce samples in a tube containing growth media and ball bearings to macerate the tissue and release contaminating pathogens followed by an overnight incubation at 41.5°C. Pathogen cell lysis and DNA release is achieved by incubating the enriched sample at 95°C for 30 minutes. LAMP DNA amplification is performed on a diluted sample of the overnight culture by incubating the reaction in a water bath using a Styrofoam box containing 63°C water. The results of the DNA amplification are visualized by adding flocculation solution. In the absence of amplification the solution will remain black and non-transparent (upper tube), alternatively if the pathogen is detected, the particles in the solution will flocculate and rapidly settle on the bottom of the tube (lower tube).

**Figure 5.**
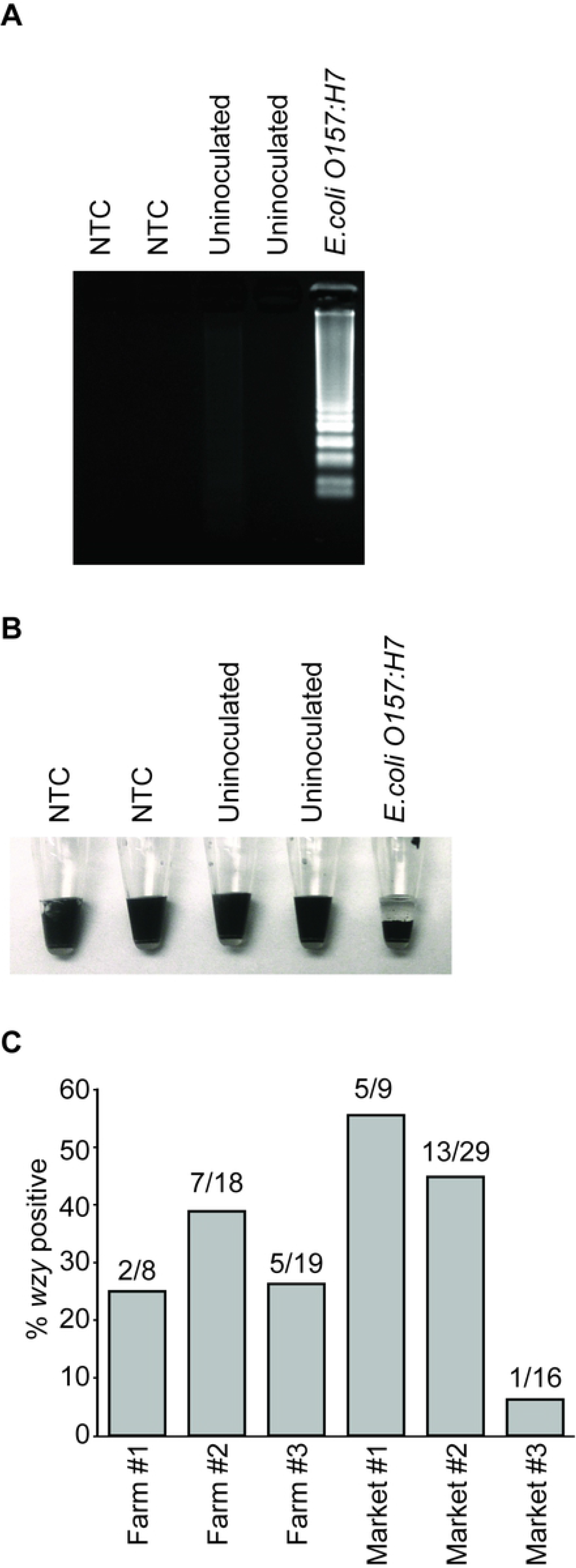
Evaluation of the complete *E. coli O157:H7* diagnostic system. (A) Freeze dried LAMP reactions containing the *wzy-1* primer set were rehydrated in betaine solution and 1 μl of diluted enriched culture from Brussel sprout leaf samples inoculated with *E. coli O157:H7*. Non-inoculated leaf cultures were used as negative controls as well as water (NTC). (B) 20 μl of flocculation solution was added to the above LAMP reactions and the solutions mixed for 10-15 seconds. Settling of the particles indicates a positive reaction. (C) Fresh produce was collected from three farms and three markets in Cambodia. Samples were assayed for the presence *wzy* using the developed diagnostic workflow. Data shows the percentage of positive samples (bar graphs) as well as the number of positive and total number of samples from each sampling site (numbers above each bar).

To assess the workflow in field conditions we transported enough reactions to test 100 vegetable samples to Cambodia with the *wzy-1* primer set. Cauliflower, radish and bean leaves, as well as bean pods were collected from three different Cambodian farms (S2 Fig) located within an approximate 10km radius of each other. Tissues were processed following the above described workflow (Fig. 4) and positive tests were obtained at all sites. In total, 25%, 39% and 26% of samples tested positive for *wzy* on farms #1, #2 and #3, respectively.

We also collected samples from three different outlets: a roadside vegetable stall and two large food markets located in a rural area (Figs S3A and S3B) and within the city Phnom Penh (Figs S3C and S3D). The roadside vegetable stall (market #1) was located along a dirt road that was used by local farmers and villagers as well as their farm animals. We found that 56% of the samples obtained from this stall tested positive (Fig 5C, Market #1). Market #2 was located in a small rural village and had a hard packed dirt floor containing numerous puddles of liquid and a low tin roof that protected the produce from direct sun exposure but generated an intense heat with little air flow (market #2, Figs S3A and S3B). Raw meat products were not refrigerated and were allowed to come in direct contact with fresh produce (Fig S3B) in closely packed stalls where a large number of flies freely moved between the meat and fresh produce. A high proportion (45%) of the samples collected from this market tested positive for *wzy* (Fig 5C). In contrast, the market located within the city (Phnom Penh) (market #3, Figs S3C and S3D) had significantly better ventilation than the rural market and a large proportion of the floor was concreted and appeared clean. The vegetables in this market looked fresh and not wilted like in the rural market. Only one out of 16 samples (6%) from this market tested positive using the *wzy* primer set (Fig 5C, Market #3).

## Discussion

The goal of this study was to develop a complete diagnostic workflow for food pathogens tailored to countries with limited resources to enhance food biosecurity capability. Our focus was to create a robust, simple and low-cost system that is safe to perform by people with limited training and equipment. The successful testing of our system in Cambodia on samples collected from the local farms and markets, suggests that we have developed a practical system that meets these requirements and is capable of providing meaningful data that can support food safety initiatives.

Culture-based techniques are the most reliable method for food pathogen identification however, these techniques are slow and require highly skilled technicians to perform [33]. The diagnostic workflow presented here is simpler and faster allowing large numbers of samples to be screened in situations where access to suitability trained microbiologists is limited. Our diagnostic workflow takes advantage of the many desirable characteristics of LAMP amplification, including high specificity and the ability to be performed in a simple water bath [34, 35]. However, like all DNA amplification reactions, LAMP is not immune to issues of contamination or false-positive amplifications [36]. Thus, it is critically important to optimize sample processing to prevent cross-contamination and develop robust specific primer sets for each target organisms prior to deployment.

The diagnostic workflow developed here can be easily adapted to detect the presence of target genes from different food pathogens using highly specific LAMP primer sets readily available in the literature [37–39]. Unlike specificity, the sensitivity of the primers is less critical to the success of the assay as the overnight enrichment step significantly increases the pathogen levels in the tested sample. Consistent with this, our assay has successfully detected the presence of 1 CFU of *E. coli O157* on a 1 g sample of Alfalfa sprouts (Fig 2B) emphasizing that the simplicity of the detection workflow presented here does not limit its capacity to detect trace amounts of pathogens on produce.

The focus of our investigation in Cambodia was to examine the performance of our workflow rather than perform a comprehensive survey of Cambodian produce. Thus, we purposely biased the sampling by seeking out produce with increased likelihood of *E. coli* contamination such as containing mud splashes or selecting farms with free roaming animals [10, 23, 40, 41]. The data obtained in this study revealed a clear correlation between the origin and condition of the produce and the likelihood of contamination. For example, the location of the food stall (market #1) on a dirt road used by local villagers and their farm animals allowed dirt from the road, potentially carrying zoonotic diseases from animal waste, to blow over the stall increasing the risk of contamination (Fig 5C) [42, 43]. Similarly, the poor drainage, problems with flies and lack of separation between raw meat and fresh produce by some vendors in market #2 also increased the risk of zoonotic disease contamination in fresh produce. Therefore, it is not surprising that these rural locations (market #1 and #2) accounted for 95% of the *wzy*-positive market samples. Consistent with this, other studies that have shown a strong correlation between food-borne disease infections and the selling of produce in markets where the vendors do not follow safe food handling practices or lack access to essential facilities (e.g. clean water, garbage removal) [3, 42]. Furthermore, a study by the World Health Organization of ‘Morning glory’ produce harvested from areas surrounding Phnom Penh, found that 100% of samples collected in Cambodian markets were contaminated with *E. coli* species [44]. Collectively, these findings suggest that there is a real need for simple, low-cost diagnostic systems to help the local authorities to improve food safety.

All the surveyed farms had a similar proportion (25-38%) of produce that tested positive for *wzy* (Fig 5C). As the farms were located within a 10 km radius of each other, these findings suggest that they are exposed to similar levels of pathogenic bacteria through common water supplies [40] or airborne particulates that can move between farms [45]. In this proof-of-concept study, we demonstrate that our simple workflow is capable of obtaining meaningful data on the prevalence of harmful food pathogens in specific locations; such information is critical to improve regional food safety.

There are many diagnostic systems for food pathogens described in the scientific literature or commercially available. However, our diagnostic workflow has a number of advantages over many of the available systems. Compared to some commercially available diagnostic systems (e.g. lateral flow strips ($6.80 USD each, Romer labs)) or systems that involve disposable electronics or custom-made microfluidic parts [46, 47]; our diagnostic system is considerably more affordable; costing $0.76 USD, including all tubes and reagents (S2 Table), which can be further reduced to $0.53 USD if the ball bearings and 50ml tubes are carefully decontaminated and reused. Cost is a critical factor for developing countries with very limited budgets devoted to food safety. Lowering the cost of assays will allow increased testing and thereby boosting its efficiency as a biosecurity tool [48].

The use of LAMP, an isothermal amplification method, allowed us to avoid the expensive thermocyclers needed in conventional DNA amplification techniques such as real-time PCR, substituting it with a simple Styrofoam box. Water was easily adjusted to the desired temperature (63°C) by mixing hot and cold water and using a standard thermometer. A lid was placed on top of the Styrofoam box to keep the water close to the initial temperature for the time needed to perform the reaction. Using this system, the water temperature dropped only 3°C over the 50-minute incubation period. This easily accessible and low-tech approach eliminates the need for expensive scientific equipment for DNA amplification, and with well-designed LAMP primers, can provide accurate detection of pathogens from food samples [21, 49].

The flocculation-based readout used in our workflow is cheap and does not require any specialized equipment, unlike conventional techniques like agarose gel electrophoresis or modern microfluidic or electronic-based systems. The flocculation solution causes the large DNA amplicons produced in the LAMP reaction to bind to suspended black charcoal particles, forming large clumps that rapidly drop to the bottom of the tube leaving a clear upper phase that is easy to distinguish from the black, non-transparent negative reactions [20]. The simplicity of the flocculation assay enables people with limited scientific training to perform and interpret the assay.

The ability to perform sample maceration, enrichment and DNA extraction in a single tube without needing to open it between steps makes processing the samples simple and safe and allows the user to handle a large number of samples at once. This process could be made even simpler by using the newly developed 30 second dipstick-based DNA purification technology [50] circumventing the need for any pipetting. Unfortunately, the rapid dipstick purification method was developed after the completion of this study.

In summary, we have developed a complete food pathogen diagnostic system composed of four key steps: pathogen enrichment, cell lysis/DNA release, LAMP amplification, and naked eye readout (Fig 4). The minimal number of steps and equipment involved, as well as the simplicity of the presence/absence flocculation readout, allows almost anyone, including those with limited scientific training, to perform the assay. Furthermore, the low cost of the system ($0.76 USD) and broad availability of its reagents, makes the system accessible to countries or institutions with limited resources who might not otherwise be able to afford regular food testing. Every step in the diagnostic workflow has been designed with the World Health Organization’s ASSURED philosophy in mind, that is, Affordable, Sensitive, Specific, User-friendly, Rapid, Equipment-free, and Deliverable to those who need it [51]. As the system is based on DNA amplification using specific primers, our diagnostic system can be easily modified to identify a large variety of pathogens including those that infect humans, crops or animals. Therefore, we anticipate that the incorporation of our diagnostic system into food safety programs of developing countries will facilitate improvements to both their food quality and human health.

## Methods

### Bacterial strains

*Escherichia coli O157:H7* str. EDL933 and *Salmonella enterica* serovar *enteritidis* were used to develop and test the diagnostic assay.

### Primer selection

A number of potential *E. coli O157:H7* target genes were selected for primer development including the shiga toxin genes *stx1* and *stx2*, the O-antigen polymerase *wzy*, the O-antigen transporter, and *rfbE*, the H7 flagellar antigen *FliC* (S1 Table). A CLUSTALW alignment was performed for each of each of these genes using *E. coli O157:H7* sequences found on the Genbank database. These alignments were used to design LAMP primer against conserved regions of these target genes using Primer Explorer software V4 (http://primerexplorer.jp/e/).

### LAMP DNA amplification

Unless otherwise stated, LAMP reactions were performed by in a solution containing 20 mM Tris (pH 8.8), 10 mM (NH_4_)_2_SO_4_, 50 mM KCl, 0.1% (v/v) Tween-20, 0.8 M betaine, 8 mM MgSO_4_, 1.2 mM dNTPs, 0.32 U/μl Bst2.0 warm-start (NEB Biolabs, USA), 0.8 μM of FIP and BIP primers and 0.2 μM of F3 and B3 primers. Reactions were incubated at 63°C for 50 minutes followed by a five-minute incubation at 80°C to denature the enzyme.

### Flocculation solution preparation

The final, optimized flocculation solution is made from 100-400 mesh activated charcoal (Sigma, St. Louis, USA) and powdered diatomaceous earth that had been ground separately in a coffee grinder for 45 seconds to break up any large particles. 400mg of activated charcoal and 600mg of diatomaceous earth were combined in a 50 ml solution containing 50 mM Tris (pH 8), 10 mM spermine and 1% (w/v) PEG8000. The flocculation solution can either be stored at 4°C or −20°C for at least a year without loss of activity.

### Freeze drying method

The buffer of Bst 2.0 warm-start DNA polymerase (NEB) was replaced with Isothermal amplification buffer (NEB) by centrifugation of the enzyme in an Amicon Ultracel-30 centrifugal filter device (Merck-Milipore) and subsequently washing in Isothermal amplification buffer twice. Aliquot 3.49 μl of LAMP solution (1.1 U/μl of dialyzed Bst 2.0 warm-start polymerase, 50 mM Tris (pH 8.8), 25 mM (NH_4_)_2_SO_4_, 25 mM KCl, 20 mM MgSO_4_, 0.25% (v/v) Triton x100, 3.4 mM dNTPs, 4.6 μM FIP and BIP primers, 0.6 μM B3 and F3 primers, 5% (w/v) Trehalose) into individual 0.2 ml tubes. The solutions were frozen in liquid nitrogen and then immediately transferred to a freeze drier for at least 4 hours under a vacuum of below 150 mTorr.

### Vegetable sample collection

In May, 2015, vegetable samples were collected from three Cambodian farms and four markets either in or surrounding Phnom Penh. The markets included a small rural village roadside vegetable stall, two large markets in the outskirts of Phnom Penh and one market within the city. Vegetables collected included cauliflower leaves, beans, sprouts, lettuce and other leafy herbs. Vegetable samples were immediately placed in individual plastic bags that were labelled and stored on ice until they were processed using the complete food pathogen diagnostic system detailed below. Access to farms was arranged by the Cambodian General Directorate of Agriculture (GDA) who obtained verbal approval from the landowners for us to collect plant samples. No protected species were samples in this study.

### Complete food pathogen diagnostic system

Approximately 1g of vegetable leaf material was placed in a 50 ml tube containing three ball bearings and 10 ml of buffered peptone growth media. The buffered peptone media was pre-aliquoted powder and made up with boiled water immediately prior to use. The tube containing the vegetable sample was sealed with Parafilm M (Bemis, WI, USA) and then shaken vigorously for one minute to aid the release of any food pathogens from the tissue. The samples were incubated overnight at 41.5°C overnight to enrich the bacteria present and then boiled for 30 minutes to both kill any bacteria present and to release their DNA into the media. One microliter of culture that had been diluted 15-fold with water was added to a freeze-dried LAMP reaction that had been rehydrate with 9 μl 0.89M betaine. The reaction was incubated in a lidded Styrofoam box containing 63°C water for 50 minutes. 20 μl of flocculation solution (50 mM Tris (pH 8), 10 mM spermine, 1% (w/v) PEG 8000, 0.8% (w/v) powdered activated charcoal, 1.2% (w/v) powdered diatomaceous earth) was then added to each LAMP reaction and the tube gently flicked to encourage mixing of the solutions. Samples in which the particles flocculated and settled on the bottom of the tube within 20 seconds were called positive, whereas those in which the particles remained suspended were negative.

## Acknowledgements

We would like to thank the many people from the General Directorate of Agriculture (GDA) in Cambodia who provided us with invaluable assistance by guiding us to different farms and markets in and around Phnom Penh. We would also like to thank Australian Centre for International Agricultural Research (ACIAR) for funding this research.

## Supporting information captions

**S1 Figure.**
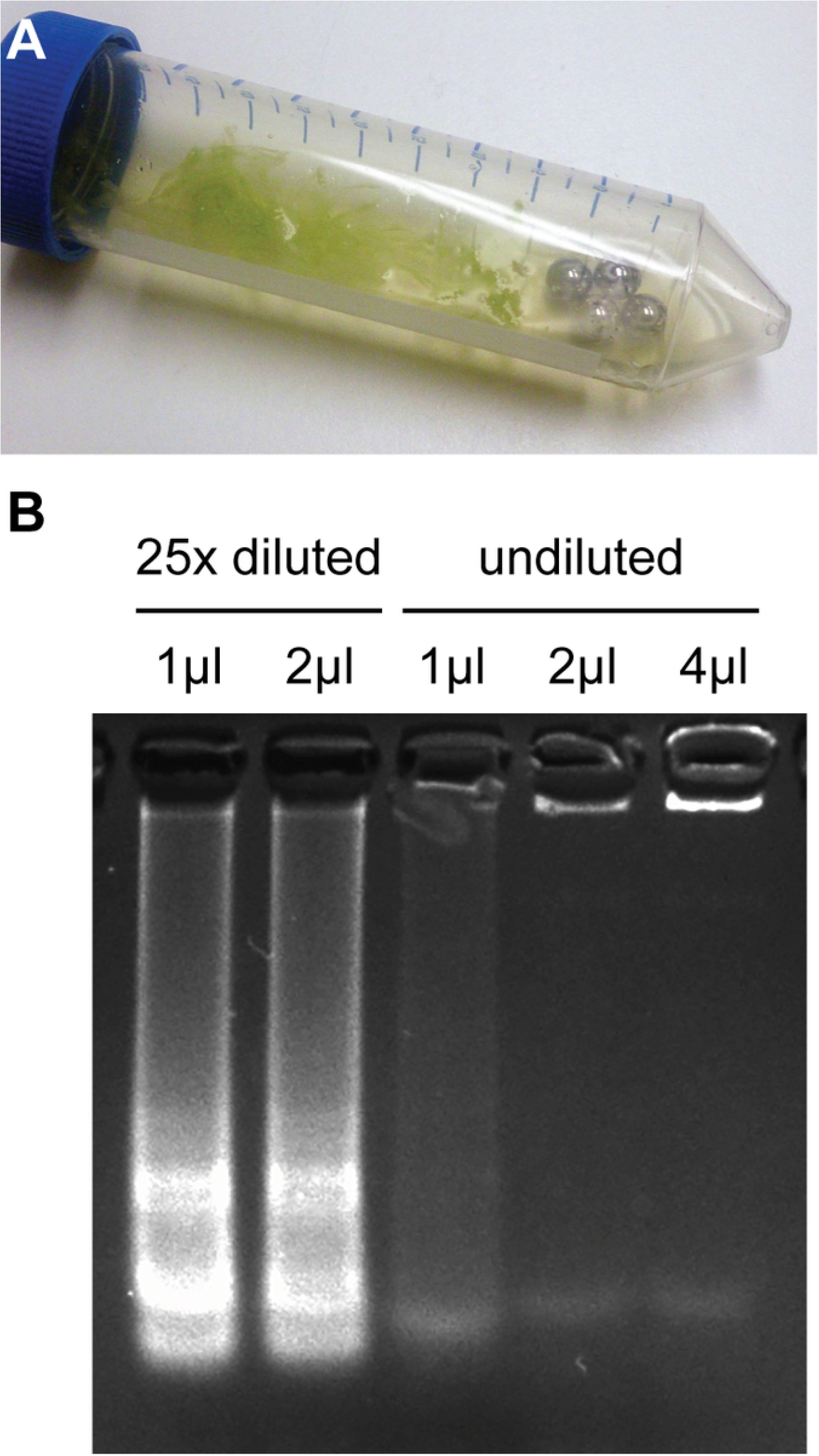
Optimization of Pathogen DNA extraction. (A) Image shows tissue that has been macerated after shaking in a 50 mL tube containing 10 ml of buffered peptone water and four ball bearings. (B) LAMP amplification products obtained using *stx2-1* primer set and template consisting of either 1, 2 or 4 μl of undiluted enriched culture or 1 or 2 μl of culture diluted 25-fold in water.

**S2 Figure.**
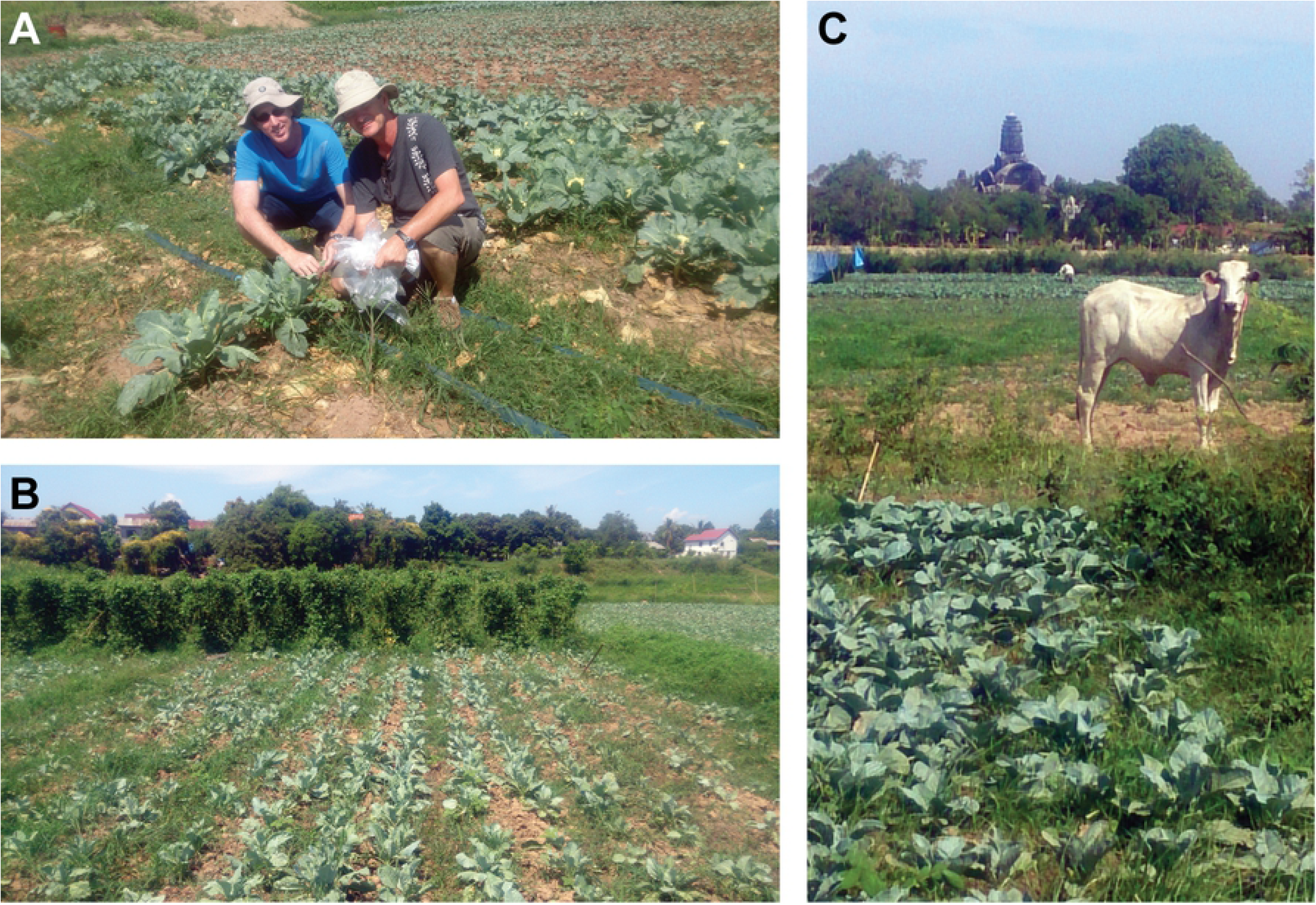
Images of Cambodian farms collection sites. (A) Collection of cauliflower leaves in individual plastic bags at farm #1. (B) Image of farm #2 which was a small plot in with cauliflower grown in the foreground and beans growing in the background. (C) Image of a cow that freely roamed amongst the cauliflower crop at farm #1.

**S3 Figure.**
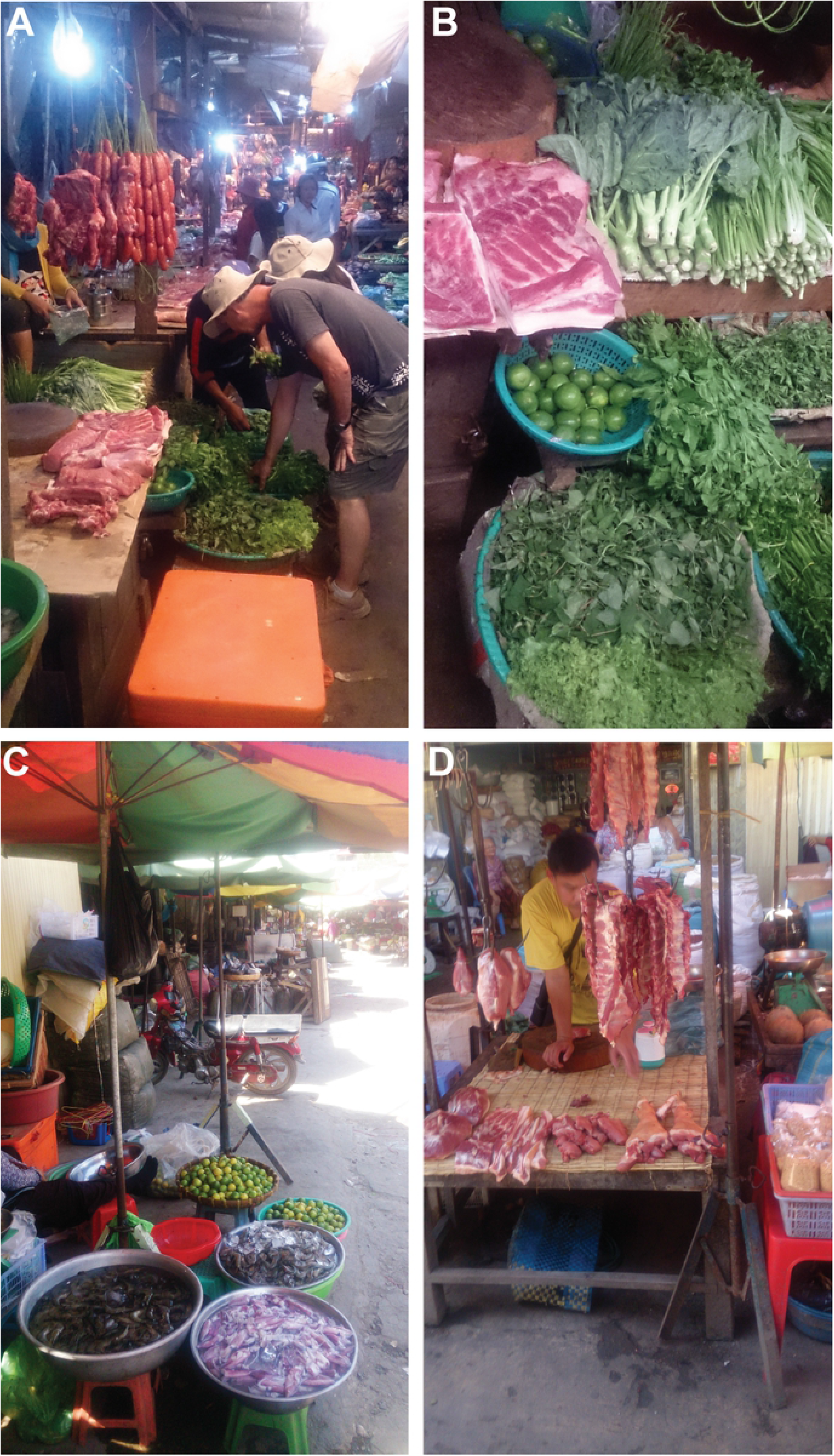
Images of Cambodian market collection sites. (A) Image of the cramped conditions in the rural market #2. (B) Image of some of the meat and produce being sold side by side at market #2. (C) Image of a large city market in Phenom Penh (market #3) with material covered tents and less cramped conditions compared to the rural market. (D) Separation of meat and fresh produce at market #3.

**S1 Table. LAMP primer sequences used in this study**

**S2 Table. Costings of individual components of the food pathogen assay.**

